# Diagnostic Performance of Fluorescence-Based Rapid On-Site Specimen Evaluation for *Helicobacter pylori* Antimicrobial Susceptibility Testing Before and After Algorithm Optimization: A Two-Round Comparative Study

**DOI:** 10.64898/2026.07.23.740309

**Authors:** Binghui Li, Lin Zhang, Yanhong Hou, Kai Wu, Jiafeng Han, Junyu Liu, Jing Zhang, Mi Yang

**Affiliations:** Department of Gastroenterology, The Eighth Medical Center of Chinese PLA General Hospital, Beijing 100091, China

**Author notes:** Corresponding author: Lin Zhang, Department of Gastroenterology, The Eighth Medical Center of Chinese PLA General Hospital, Beijing 100091, China.

**Keywords:** *Helicobacter pylori*, antimicrobial susceptibility testing, fluorescence-based rapid on-site specimen evaluation, algorithm optimization, diagnostic performance

## Abstract

**Objective:** This study was designed to evaluate the diagnostic performance of fluorescence-based rapid on-site specimen evaluation (F-ROSE) for antimicrobial susceptibility testing of *Helicobacter pylori* (*H. pylori*) and to compare test performance before and after iterative optimization of fluorescence image acquisition and of the artificial intelligence (AI) recognition algorithm, using two consecutive rounds of paired testing, so as to provide trial data supporting clinical implementation of rapid susceptibility testing.

**Methods:** Forty patients with a urea breath test (UBT) or rapid urease test (RUT) positive for *H. pylori* within the preceding 2 weeks, together with strongly positive endoscopic findings, were prospectively enrolled. In order of enrollment they were allocated to two rounds in which F-ROSE was compared with culture-based susceptibility testing. Twenty patients were tested in round 1 on an automated fluorescence immunoassay scanner running the original algorithm; a further 20 were tested in round 2 with the optimized algorithm. With in vitro culture and E-test as the reference standard, sensitivity, specificity, positive predictive value (PPV), negative predictive value (NPV) and accuracy of F-ROSE were calculated for each round for amoxicillin, levofloxacin hydrochloride and clarithromycin, and agreement between methods was assessed with Cohen’s kappa. Receiver operating characteristic (ROC) analysis was used to identify the optimal cutoff of the fluorescence residual rate (FRR) and to determine how far threshold adjustment improved performance.

**Results:** Culture succeeded in 15 patients in round 1 and in 14 patients in round 2, leaving 29 evaluable samples. Overall diagnostic performance after algorithm optimization was clearly better than before. Pooled across the three antibiotics, sensitivity was 92.9%, specificity 71.0% and accuracy 77.8% before optimization; after optimization sensitivity rose to 100.0%, specificity increased slightly to 74.2% and accuracy to 81.0%. NPV reached 100.0% for all three agents after optimization, and no resistant isolate was missed. The gain was largest for clarithromycin, for which sensitivity rose from 85.7% to 100.0%, accuracy from 66.7% to 78.6% and kappa from 0.348 to 0.571, indicating a clear improvement in agreement between the two methods. ROC analysis showed that drug-specific optimal cutoffs derived from the Youden index improved specificity appreciably compared with the uniform 50% FRR threshold currently applied, the gain being most evident for levofloxacin hydrochloride.

**Conclusions:** With sharper fluorescence images and an improved AI recognition algorithm, F-ROSE performs better in *H. pylori* susceptibility testing, and the reduction in missed resistant strains is particularly noteworthy; the assay is therefore a plausible option for rapid susceptibility screening in clinical practice. Multicenter studies with larger samples are still required to confirm the stability of the technique.

## 1. Introduction

*Helicobacter pylori* (*H. pylori*) is a gram-negative, microaerophilic organism that colonizes the surface epithelium of the human gastric mucosa, and more than half of the world population is infected^[1]^. The organism is closely associated with chronic gastritis, peptic ulcer disease, gastric mucosa-associated lymphoid tissue (MALT) lymphoma and gastric cancer, and the International Agency for Research on Cancer of the World Health Organization has classified it as a group I carcinogen^[2]^. Eradication of *H. pylori* is therefore a key step in reducing this disease burden^[3]^. Wide use of antibiotics in recent years has nevertheless made resistance an increasingly serious problem. Multicenter data from China show resistance rates exceeding 30% for clarithromycin and 60% for metronidazole, with levofloxacin resistance also on the rise^[4]^. Where resistance is this common, eradication rates achieved with empirical regimens fall markedly, and individualized susceptibility testing to guide precise antibiotic selection has become an urgent clinical need^[5]^.

The current reference standard for H. pylori susceptibility testing is in vitro culture combined with agar dilution or E-test. This phenotypic method has high accuracy and reproducibility, yet it has prominent limitations: culture incubation takes one to two weeks, strict requirements are required for specimen transportation and microaerobic incubation, and bacterial isolation success rates vary widely across clinical laboratories, ranging from 60% to 80% in routine clinical practice, accompanied by cumbersome experimental workflows^[6]^. None of this meets the clinical demand for a rapid susceptibility result. A method that can deliver reliable susceptibility data within a short time would therefore be of considerable practical value.

Originally developed for cytological rapid onsite diagnosis, fluorescence-based rapid on-site specimen evaluation (F-ROSE) has been translated to microbiology via improved fluorescent staining and AI image analysis^[7]^. Gastric mucosal samples stained with specific fluorescent probes are scanned automatically; AI quantifies bacterial fluorescence intensity differences between antibiotic-exposed and drug-free groups, and susceptibility is judged by fluorescence residual rate (FRR: ≤50% susceptible, >50% resistant). This assay shortens susceptibility testing from days to hours to guide personalized therapy. As an emerging technique, F-ROSE needs systematic validation, as its initial AI algorithm suffers interference from uneven image quality and heterogeneous Helicobacter pylori morphology. Iterative optimization of imaging and bacterial recognition may boost its diagnostic accuracy. We therefore performed two parallel test rounds against culture (reference standard) before and after algorithm optimization to assess F-ROSE’s practicability for H. pylori susceptibility testing, quantify performance gains from algorithm upgrading, and supply data supporting further technical improvement and clinical application.

## 2. Materials and Methods

### 2.1 Ethics statement

Written informed consent was obtained from all participants. The protocol was reviewed and approved by the Ethics Committee of the Eighth Medical Center of Chinese PLA General Hospital (approval number S-2026-003-01) and registered with the Chinese Clinical Trial Registry (http://www.chictr.org.cn/) on 21 May 2026 under registration number ChiCTR2600125137.

### 2.2 Study population

This was a single-center, prospective, sequentially grouped diagnostic study. Between January and June 2026, 40 adult patients with strongly positive endoscopic findings for *H. pylori* were prospectively enrolled at the Eighth Medical Center of Chinese PLA General Hospital. Inclusion criteria were: (1) *H. pylori* positivity confirmed by UBT or RUT within 2 weeks before enrollment; (2) no history of *H. pylori* eradication therapy and no systemic antibiotic exposure of any kind (including exposure unrelated to eradication) within 4 weeks; and (3) no use of proton pump inhibitors within 2 weeks.

The 40 patients were assigned in chronological order to two rounds. In round 1 (before algorithm optimization) 20 patients were enrolled, of whom 15 yielded a successful culture and were included in the final analysis; 5 cultures failed and were excluded. In round 2 (after algorithm optimization) 20 patients were enrolled, of whom 14 yielded a successful culture and entered the analysis, while 6 cultures failed and were excluded. All 40 specimens tested positive for Hp by F-ROSE, and susceptibility results for all three antibiotics were obtained in every case. Culture failure was attributable mainly to bacterial death caused by delayed transport, insufficient bacterial load in the specimen and contamination by commensal flora. The 29 culture-positive specimens from the two rounds were pooled for the subsequent analysis of diagnostic performance.

### 2.3 Test methods

#### 2.3.1 Culture (reference standard)

One antral mucosal biopsy specimen was taken during endoscopy, placed in a sterile tube containing dedicated transport medium and delivered to the microbiology laboratory within 4 h. After thorough homogenization the specimen was inoculated onto Columbia blood agar containing selective antimicrobial supplements and incubated at 37 °C under microaerophilic conditions (5% O□, 10% CO□, 85% N□) for 3–5 days. When suspicious colonies appeared, *H. pylori* was identified on the basis of colony morphology, Gram staining, and urease, catalase and oxidase tests. Minimum inhibitory concentrations (MICs) of the three first-line antibiotics amoxicillin, levofloxacin hydrochloride and clarithromycin were determined by E-test, and breakpoints followed the European Committee on Antimicrobial Susceptibility Testing (EUCAST) MIC criteria for *H. pylori*: amoxicillin ≤0.125 μg/mL susceptible and >0.125 μg/mL resistant; clarithromycin ≤0.125 μg/mL susceptible and >0.5 μg/mL resistant; and levofloxacin ≤1.0 μg/mL susceptible and >1.0 μg/mL resistant^[8]^.

#### 2.3.2 F-ROSE

One antral mucosal biopsy specimen was obtained endoscopically and placed immediately in 1 mL of sterile phosphate-buffered saline (PBS), then ground at a steady rate with a sterile pestle for 30 s to release the bacteria colonizing the mucosa and to produce a bacterial homogenate. The homogenate was divided into four 200 μL aliquots, which were dispensed into wells containing amoxicillin, levofloxacin hydrochloride or clarithromycin and into an antibiotic-free control well. After incubation at 37 °C for 1 h, 100 μL of F-ROSE reagent (a proprietary formulation containing the two specific fluorescent probes EthD-III and DMAO, which label dead and live bacteria respectively) was added to each well, and the plate was incubated for 5 min at room temperature in the dark. After incubation, fluorescence images of all wells are captured by an automated fluorescence immuno-scanner within a 15-minute scanning procedure, leading to a total assay runtime of approximately 1–2 hours. An AI algorithm performed automated bacterial recognition, classification and quantitative counting on the fluorescence images and calculated the fluorescence residual rate (FRR), which was used to assess resistance. FRR was calculated as:

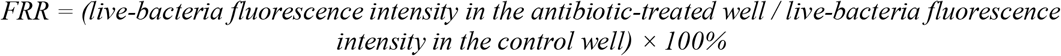

Susceptibility was interpreted against a 50% FRR threshold: an FRR ≤ 50% was read as susceptible (S), that is, the antibiotic effectively suppressed bacterial viability, whereas an FRR > 50% was read as resistant (R), indicating a markedly reduced response to the agent concerned.

### 2.4 Content of the algorithm optimization

After the first round had been completed, the AI analysis algorithm of the F-ROSE system was upgraded in two respects. The first concerned fluorescence image acquisition and preprocessing: excitation parameters were retuned and the preprocessing algorithm refined, which reduced background noise, raised the signal-to-noise ratio of the fluorescence signal and improved image clarity and quality considerably. The second concerned the intelligent recognition model for *H. pylori*: the training set was expanded to cover more scenarios and more bacterial morphologies, and the deep learning architecture was refined iteratively, which improved recognition of atypical forms such as coccoid *H. pylori* and lowered both false and missed bacterial identification. Round 2 was performed entirely with the optimized version of the algorithm. Specimen handling, reagent lots, incubation conditions and susceptibility interpretation criteria were kept identical to those of round 1 throughout, so that data from the two rounds remained consistent and comparable.

### 2.5 Statistical analysis

Data were analyzed with SPSS 26.0. Continuous variables are expressed as mean ± standard deviation (x□ ± s), and age was compared between the two groups with the independent-samples t test. Categorical variables are described as counts (percentages); sex distribution and culture success rate were compared with the χ^2^ test or Fisher’s exact test. The significance level was set at α = 0.05, and P < 0.05 was considered statistically significant.

With culture-based susceptibility results as the reference standard, the following measures of diagnostic performance were calculated for F-ROSE: sensitivity, specificity, positive predictive value (PPV), negative predictive value (NPV) and accuracy. Sensitivity was defined as the proportion of culture-defined resistant isolates correctly identified by F-ROSE, and specificity as the proportion of culture-defined susceptible isolates correctly identified by F-ROSE. Sensitivity, specificity, PPV, NPV and their 95% confidence intervals (95% CIs) were obtained with the Wilson score interval.

Cohen’s kappa was used to assess agreement between the two methods after correction for chance; its 95% CI was derived from the bias-corrected and accelerated (BCa) bootstrap (n□□□□ = 100,000). Kappa values were interpreted as follows: κ ≤ 0.20, slight agreement; 0.21–0.40, fair; 0.41–0.60, moderate; 0.61–0.80, substantial; and κ > 0.80, almost perfect agreement^[9]^.

Taking the culture-based susceptibility result as a binary reference standard (resistant = positive, susceptible = negative) and the FRR measured by F-ROSE as a continuous diagnostic variable, receiver operating characteristic (ROC) curves were plotted separately for amoxicillin, levofloxacin hydrochloride and clarithromycin, and the area under the curve (AUC) was calculated. The 95% CI of each AUC was estimated with the BCa bootstrap (n = 100,000 resamples). The FRR value corresponding to the maximum Youden index (Youden index = sensitivity + specificity − 1) was taken as the optimal diagnostic cutoff for each antibiotic^[10]^, and diagnostic performance at that cutoff was compared with performance at the 50% cutoff currently in use.

To assess whether F-ROSE discriminated equally well across antibiotics, pairwise comparisons of the three ROC curves were made with the nonparametric DeLong method^[11]^. This approach constructs structured statistics (placement values) from the paired samples and derives the variance–covariance matrix of the difference between the AUCs of two correlated ROC curves, yielding a Wald-type Z statistic and the corresponding two-sided P value. P < 0.05 was taken as statistically significant. The ROC analyses were based on all 29 culture-positive samples from the two rounds combined.

Analyses were performed for each antibiotic (amoxicillin, levofloxacin hydrochloride and clarithromycin) and for the three drugs pooled, and results of the two rounds were compared descriptively.

## 3. Results

### 3.1 General characteristics of the two rounds

Baseline characteristics of the two groups are shown in Table 1. Of the 20 patients in round 1, 15 were men and 5 women, aged 20–75 years, with a mean age of 46.2 ± 16.5 years; culture succeeded in 15 of the 20 specimens, a success rate of 75.0%, and the culture-positive specimens came from 12 men and 3 women. Among the 5 failed cultures, 4 plates showed no bacterial growth and 1 was discarded because of heavy contamination by commensal flora.

**Table 1.** Comparison of baseline characteristics.

| Baseline variable | Pre-optimization group<br>(n = 20) | Post-optimization<br>group (n = 20) | Statistic | P value |
| --- | --- | --- | --- | --- |
| Age (years, $\bar{x} \pm s$) | 46.2 ± 16.5 | 40.9 ± 15.4 | t = 1.071 | 0.291 |
| Age range (years) | 20–75 | 19–69 | – | – |
| Sex (male/female, n) | 15/5 | 17/3 | – | 0.695* |
| Male (%) | 75.0 | 85.0 | – | – |
| Successful culture, n (%) | 15 (75.0) | 14 (70.0) | – | 1.000* |
| Failed culture, n (%) | 5 (25.0) | 6 (30.0) | – | – |

Note: \* Sex distribution and culture success rate were compared with Fisher's exact test.

Of the 20 participants in round 2, 17 were men and 3 women, aged 19–69 years, with a mean age of 40.9 ± 15.4 years; 14 specimens grew successfully, a success rate of 70.0%, with culture-positive specimens from 12 men and 2 women. Among the 6 failures, 3 showed no colony growth and 3 were discarded for contamination.

Age (t = 1.071, P = 0.291), sex distribution (Fisher’s exact test, P = 0.695) and culture success rate (Fisher’s exact test, P = 1.000) did not differ significantly between the two groups, indicating balanced and comparable baseline data.

### 3.2 Diagnostic performance by antibiotic in round 1

Results obtained with the first-generation algorithm, set against the culture reference standard, are shown in Table 2. For amoxicillin, sensitivity was 100.0% (95% CI: 20.7%–100.0%), specificity 85.7% (95% CI: 60.1%–96.0%), PPV 33.3%, NPV 100.0% and accuracy 86.7%, with a kappa of 0.444, indicating moderate agreement between the two methods. For levofloxacin, sensitivity was likewise 100.0%, with specificity 66.7%, PPV 66.7%, NPV 100.0%, accuracy 80.0% and kappa 0.615, corresponding to substantial agreement. Clarithromycin was the only antibiotic for which the first-generation algorithm missed a resistant isolate: sensitivity was only 85.7%, with one false-negative sample; specificity was 50.0%, NPV fell to 80.0%, accuracy was 66.7% and kappa was 0.348, that is, only fair agreement. These figures point to a clear weakness of the first-generation AI model in recognizing clarithromycin-resistant strains, with the attendant risk that a resistant isolate is misclassified as susceptible.

**Table 2.** Round 1: susceptibility results obtained by F-ROSE versus culture.

| Index | Amoxicillin | Levofloxacin | Clarithromycin | Three drugs pooled |
| --- | --- | --- | --- | --- |
| TP/FP/TN/FN | 1/2/12/0 | 6/3/6/0 | 6/4/4/1 | 13/9/22/1 |
| Sensitivity | 100.0% (20.7%–100.0%) | 100.0% (61.0%–100.0%) | 85.7% (48.7%–97.4%) | 92.9% (68.5%–98.7%) |
| Specificity | 85.7% (60.1%–96.0%) | 66.7% (35.4%–87.9%) | 50.0% (21.5%–78.5%) | 71.0% (53.4%–83.9%) |
| PPV | 33.3% (6.1%–79.2%) | 66.7% (35.4%–87.9%) | 60.0% (31.3%–83.2%) | 59.1% (38.7%–76.7%) |
| NPV | 100.0% (75.7%–100.0%) | 100.0% (61.0%–100.0%) | 80.0% (37.6%–96.4%) | 95.7% (79.0%–99.2%) |
| Accuracy | 86.7% | 80.0% | 66.7% | 77.8% |
| Kappa (95% CI) | 0.444 (0.000–1.000) | 0.615 (0.237–1.000) | 0.348 (–0.125–0.737) | 0.476 (0.171–0.756) |
| Strength of agreement | Moderate | Substantial | Fair | Moderate |

Note: TP: true positive; FP: false positive; TN: true negative; FN: false negative. 95% CIs were calculated with the Wilson score interval; 95% CIs for kappa were obtained with the BCa bootstrap (n = 100,000).

### 3.3 Diagnostic performance by antibiotic in round 2

F-ROSE and culture results for round 2 are compared in Table 3. Once the imaging and recognition algorithms had been optimized, no resistant isolate was missed for any of the three antibiotics and sensitivity reached 100.0% throughout; details are given in Table 3 and Figure 1.

**Table 3.** Round 2: susceptibility results obtained by F-ROSE versus culture.

| Index | Amoxicillin | Levofloxacin | Clarithromycin | Three drugs pooled |
| --- | --- | --- | --- | --- |
| TP/FP/TN/FN | 2/2/10/0 | 2/3/9/0 | 7/3/4/0 | 11/8/23/0 |
| Sensitivity | 100.0% (34.2%–100.0%) | 100.0% (34.2%–100.0%) | 100.0% (64.6%–100.0%) | 100.0% (74.1%–100.0%) |
| Specificity | 83.3% (55.2%–95.3%) | 75.0% (46.8%–91.1%) | 57.1% (25.0%–84.2%) | 74.2% (56.8%–86.3%) |
| PPV | 50.0% (15.0%–85.0%) | 40.0% (11.8%–76.9%) | 70.0% (39.7%–89.2%) | 57.9% (36.3%–76.9%) |
| NPV | 100.0% (72.2%–100.0%) | 100.0% (70.1%–100.0%) | 100.0% (51.0%–100.0%) | 100.0% (85.7%–100.0%) |
| Accuracy | 85.7% | 78.6% | 78.6% | 81.0% |
| Kappa (95% CI) | 0.588 (–0.000–1.000) | 0.462 (0.000–1.000) | 0.571 (0.000–0.857) | 0.541 (0.250–0.791) |
| Strength of agreement | Moderate | Moderate | Moderate | Moderate |

Note: TP: true positive; FP: false positive; TN: true negative; FN: false negative. 95% CIs were calculated with the Wilson score interval; 95% CIs for kappa were obtained with the BCa bootstrap (n = 100,000).

**Figure 1.**
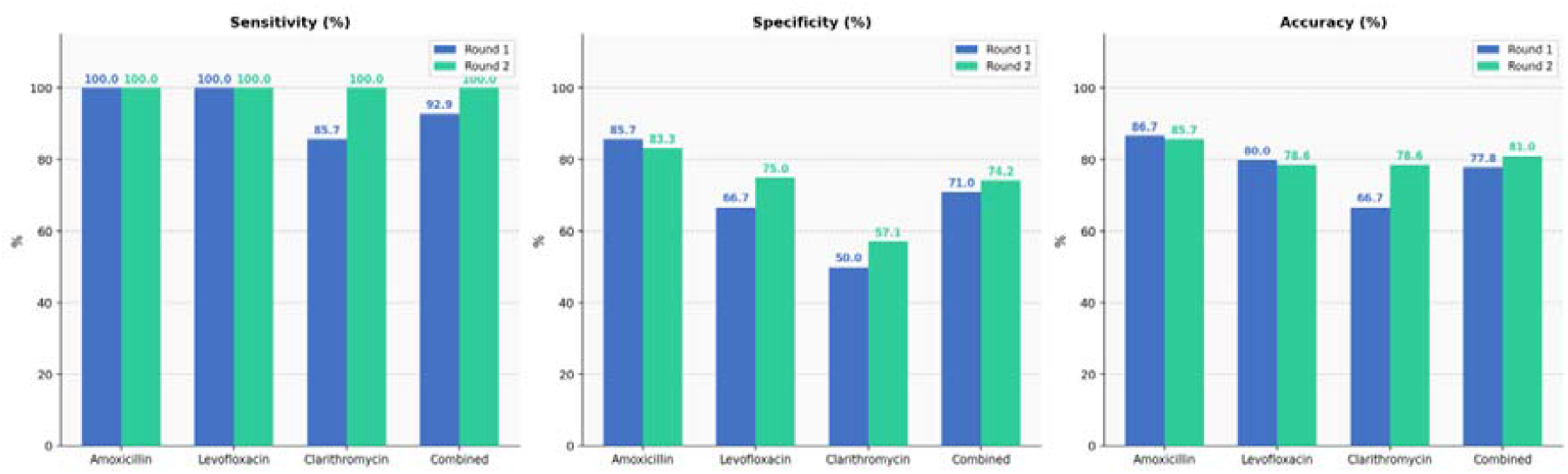
Bar chart comparing diagnostic performance between the two rounds. Note: blue bars show data from round 1 before optimization and green bars data from round 2 after algorithm optimization; the panels from left to right show sensitivity, specificity and accuracy.

Figure 1 presents sensitivity, specificity and accuracy for the two rounds as grouped bar charts. Apart from small decreases in the specificity and accuracy for amoxicillin and in the accuracy for levofloxacin, every measure improved after optimization, and the gains were most marked for clarithromycin.

The improvement for clarithromycin was the most striking, as shown in detail in Figure 2: sensitivity rose from 85.7% to 100.0% with no missed detections, accuracy from 66.7% to 78.6%, and kappa from 0.348 to 0.571, moving agreement from fair to moderate. Algorithm optimization thus improved recognition of clarithromycin-resistant strains substantially.

**Figure 2.**
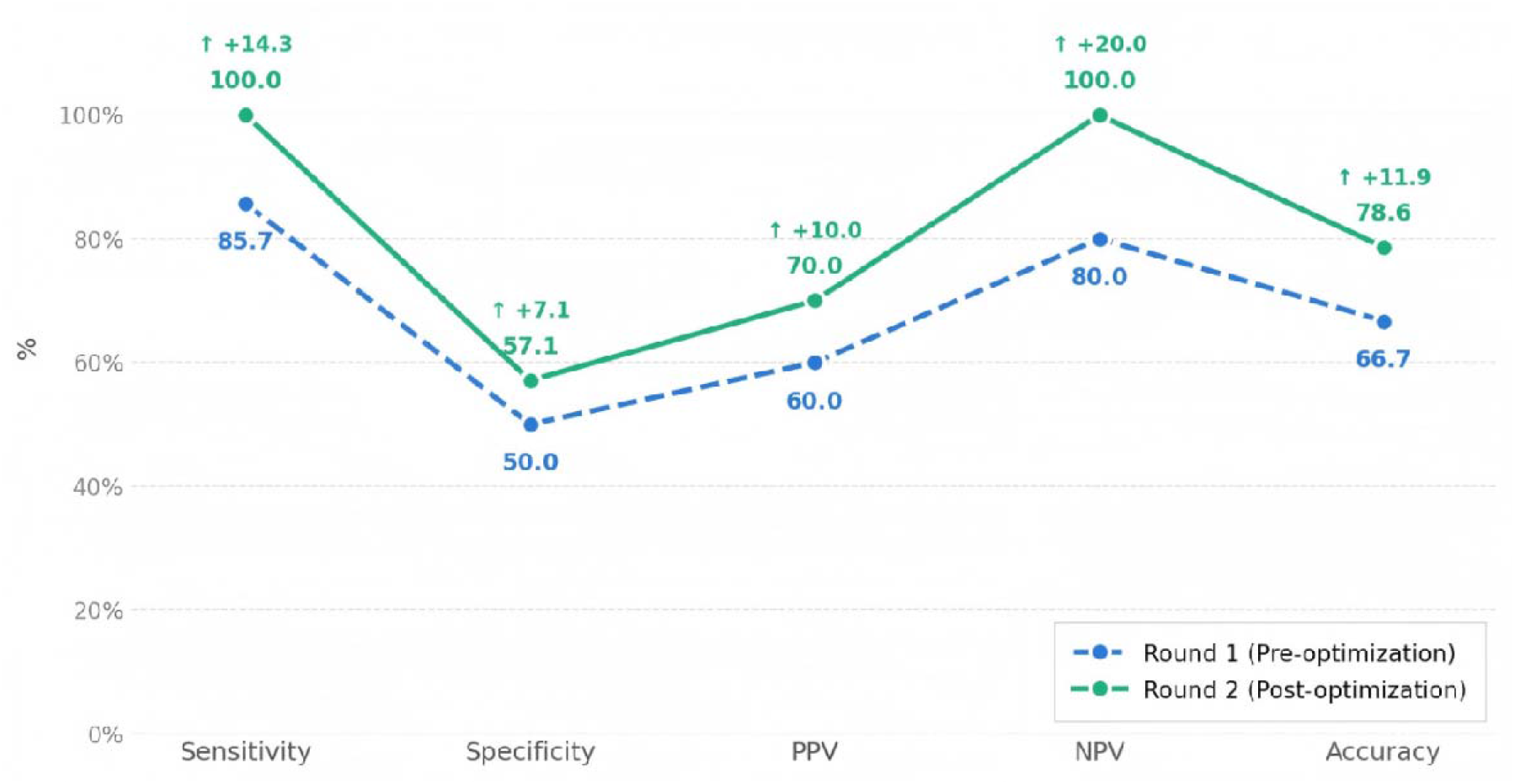
Line chart of five measures of diagnostic performance for clarithromycin before and after algorithm optimization. Note: blue dashed lines with circles denote round 1 before optimization and green solid lines with circles round 2 after optimization; the horizontal axis shows sensitivity, specificity, positive predictive value (PPV), negative predictive value (NPV) and accuracy, and the vertical axis the value of each measure as a percentage. Green upward arrows above the data points give the absolute gain achieved after optimization.

### 3.4 Combined comparison of the two rounds

When all 29 specimens and the 87 paired susceptibility results from both rounds were pooled (Table 4), overall sensitivity across the three antibiotics was 96.0%, specificity 72.6% and accuracy 79.3%, with a kappa of 0.522, indicating moderate overall agreement between the two methods.

**Table 4.** Pooled data from both rounds: susceptibility results obtained by F-ROSE versus culture.

| Index | Amoxicillin | Levofloxacin | Clarithromycin | Three drugs pooled |
| --- | --- | --- | --- | --- |
| TP/FP/TN/FN | 3/4/22/0 | 8/6/15/0 | 13/7/8/1 | 24/17/45/1 |
| Sensitivity | 100.0% (43.8%–100.0%) | 100.0% (67.6%–100.0%) | 92.9% (68.5%–98.7%) | 96.0% (80.5%–99.3%) |
| Specificity | 84.6% (66.5%–93.9%) | 71.4% (50.0%–86.2%) | 53.3% (30.1%–75.2%) | 72.6% (60.4%–82.1%) |
| PPV | 42.9% (15.8%–75.0%) | 57.1% (32.6%–78.6%) | 65.0% (43.3%–81.9%) | 58.5% (42.2%–73.2%) |
| NPV | 100.0% (85.1%–100.0%) | 100.0% (79.6%–100.0%) | 88.9% (56.5%–98.0%) | 97.8% (89.1%–99.6%) |
| Accuracy | 86.2% | 79.3% | 72.4% | 79.3% |
| Kappa (95% CI) | 0.532 (0.000–0.901) | 0.580 (0.281–0.828) | 0.455 (0.136–0.726) | 0.522 (0.299–0.720) |
| Strength of agreement | Moderate | Moderate | Moderate | Moderate |

Note: TP, true positive; FP, false positive; TN, true negative; FN, false negative. 95% CIs were calculated with the Wilson score interval; 95% CIs for kappa were obtained with the BCa bootstrap (n = 100,000).

Forest plots of sensitivity and specificity based on the Wilson score interval (Figure 3) display the single-round and pooled estimates together with their 95% CIs; circles denote round 1, squares round 2 and diamonds the pooled data, and the vertical dashed line marks 100% performance. Figure 3 shows that, with only 15 and 14 evaluable samples in rounds 1 and 2 respectively, the 95% CIs are generally wide and statistical precision is limited: although the point estimates of sensitivity for amoxicillin and levofloxacin were 100.0% in both rounds, the lower confidence limits were clearly low, and for clarithromycin in round 1 the lower limit of the sensitivity CI was only 48.7%, with an even wider interval for specificity. Intervals this wide in a single small round mean that conclusions drawn from either round alone carry considerable uncertainty and provide weak evidence. Pooling the 29 specimens from both rounds narrowed the intervals somewhat and improved statistical reliability to a degree.

**Figure 3.**
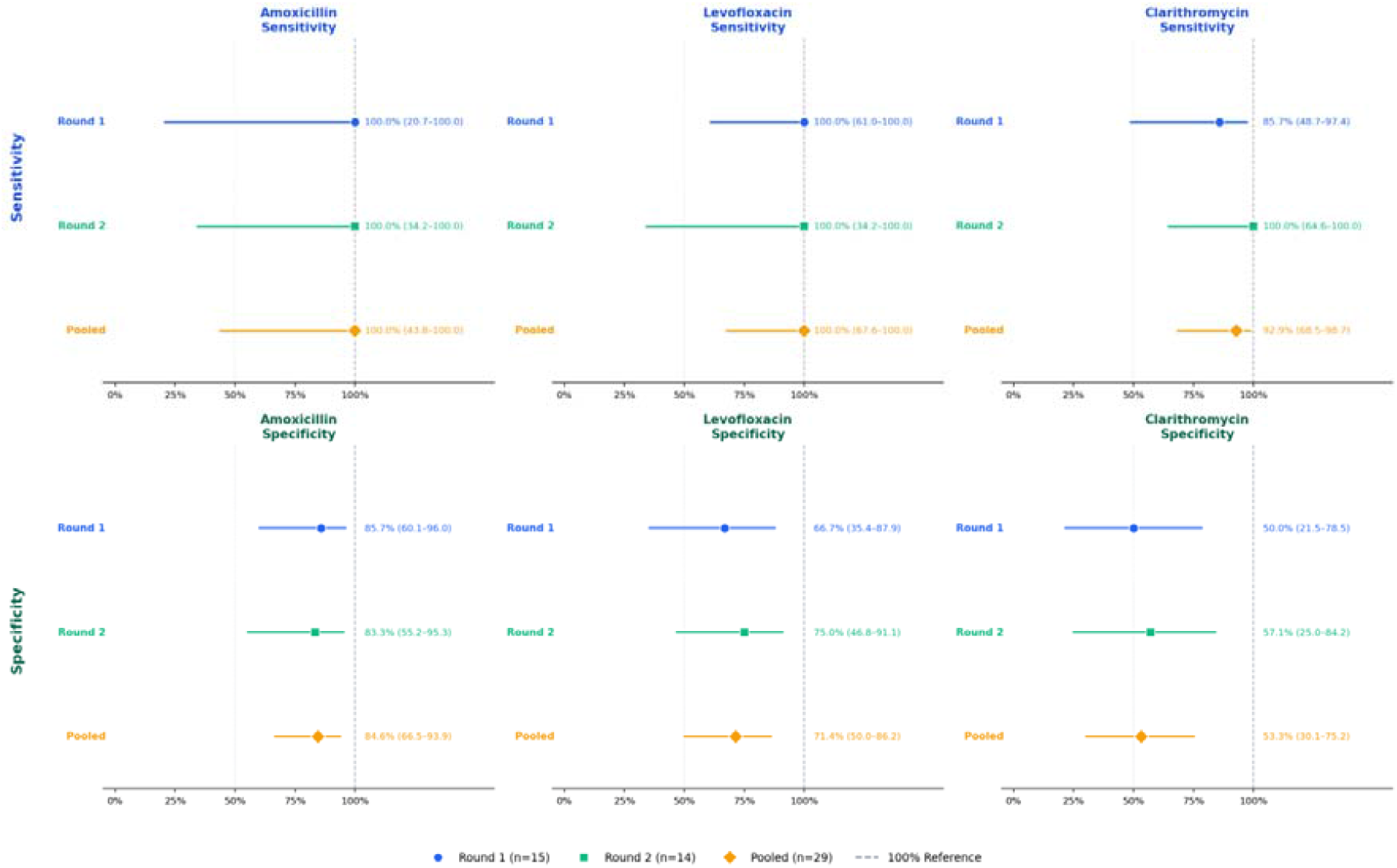
Forest plot of the diagnostic performance of F-ROSE for the three antibiotics. Note: the plot is arranged in panels, with amoxicillin, levofloxacin and clarithromycin across the three columns and sensitivity and specificity in the upper and lower blocks respectively. 95% CIs were calculated with the Wilson score interval.

### 3.5 ROC analysis and cutoff optimization

Taking the FRR values of the 29 culture-positive samples as the continuous diagnostic variable, ROC curves were plotted for each antibiotic (Figure 4). The AUC was 0.949 (95% CI: 0.821–1.000) for amoxicillin and 0.941 (95% CI: 0.826–1.000) for levofloxacin hydrochloride, both indicating good discrimination; for clarithromycin the AUC was 0.774 (95% CI: 0.581–0.931), lower than for the other two agents and with a wide confidence interval, which suggests that the stability of the estimate is limited by sample size.

**Figure 4.**
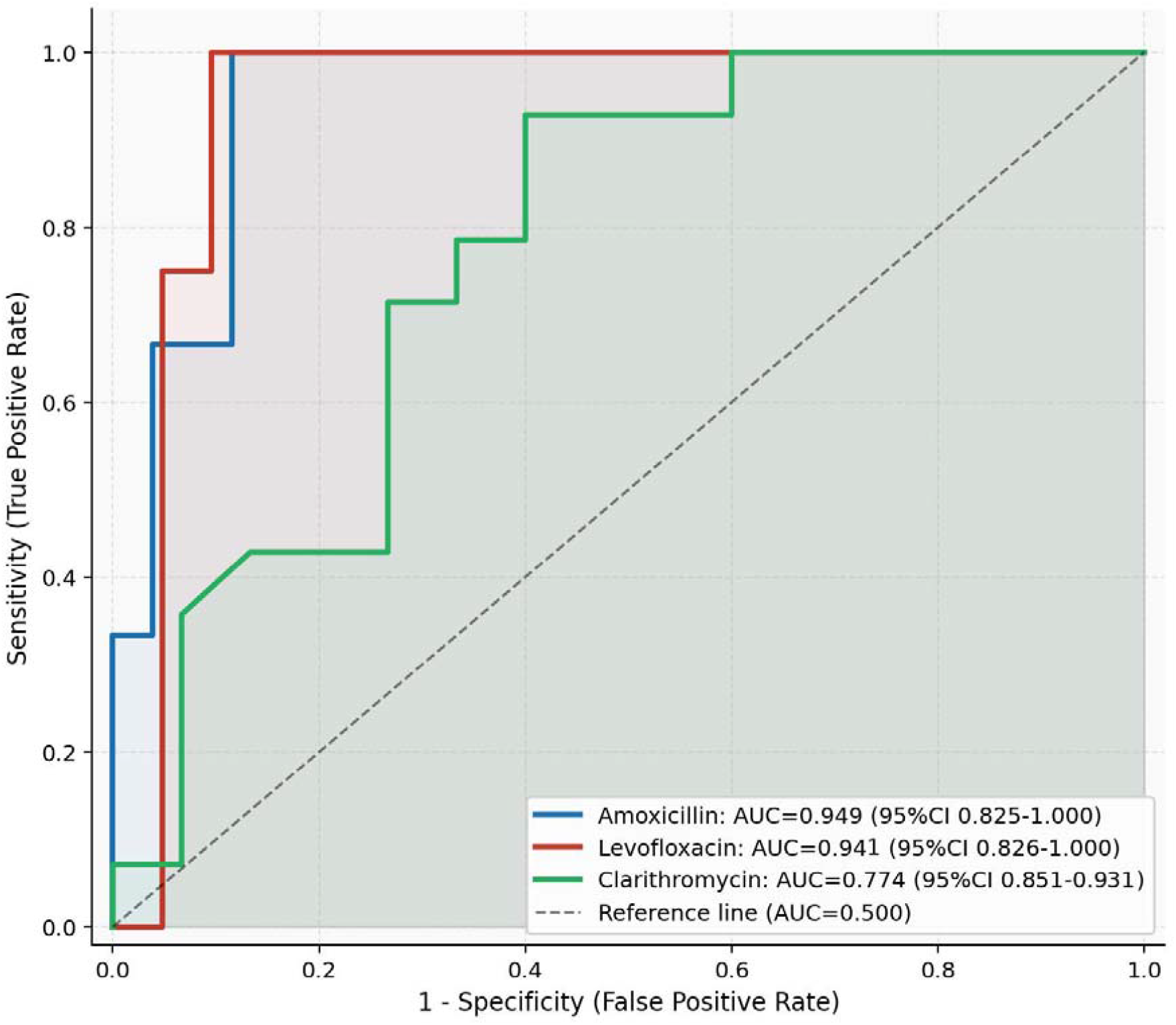
ROC curves for the three antibiotics. Note: AUC, area under the curve; its 95% CI was estimated with the bias-corrected and accelerated (BCa) bootstrap, n = 100,000. The FRR corresponding to the maximum Youden index (Youden = sensitivity + specificity − 1) was taken as the optimal diagnostic cutoff.

The optimal FRR cutoffs determined from the Youden index were 0.5556 for amoxicillin, 0.5812 for levofloxacin hydrochloride and 0.5164 for clarithromycin, all slightly above the 50% threshold currently in use. Diagnostic performance at the optimal and current cutoffs is compared in Table 5. For amoxicillin, sensitivity was 100.0% at either cutoff, while raising the threshold to 0.5556 increased specificity from 84.6% to 88.5% and removed one false positive. The improvement was clearest for levofloxacin hydrochloride: raising the cutoff from 0.5000 to 0.5812 increased specificity from 71.4% to 90.5% and PPV from 57.1% to 80.0%, reduced false positives from 6 to 2, and left sensitivity unchanged at 100.0%. For clarithromycin the cutoff shift was smaller (0.5000→0.5164) and specificity rose only modestly, from 53.3% to 60.0%, remaining low; one false negative persisted and sensitivity was 92.9%.

**Table 5.**
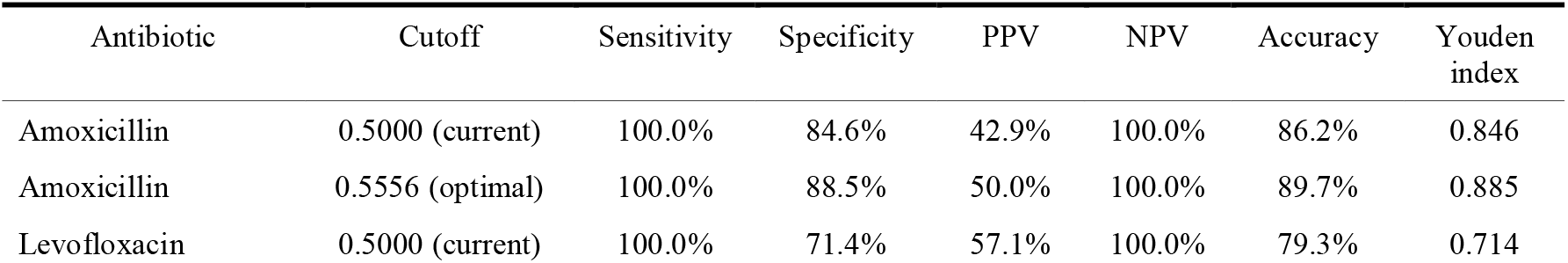
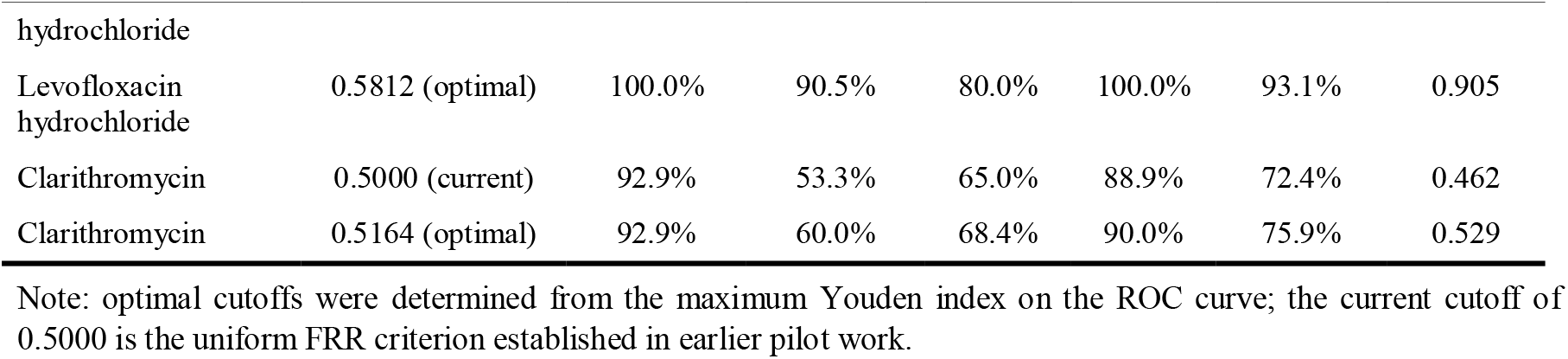
Diagnostic performance at the optimal cutoff versus the current 50% cutoff.

### 3.6 DeLong test

Pairwise comparisons of the three ROC curves by the nonparametric DeLong method are shown in Table 6. The difference in AUC between amoxicillin and levofloxacin hydrochloride was 0.008 (Z = 0.121, P = 0.904) and was not statistically significant. The difference between amoxicillin and clarithromycin was 0.175 (Z = 1.685, P = 0.092) and that between levofloxacin hydrochloride and clarithromycin 0.167 (Z = 1.603, P = 0.109), neither reaching statistical significance. Although the point estimate of the AUC for clarithromycin (0.774) was clearly lower than those for the other two agents (0.949 and 0.941), the small sample available here (only 3 amoxicillin-resistant samples) left the analysis underpowered, so the data do not establish that FRR discriminates differently across the three antibiotics.

**Table 6.** Pairwise comparison of AUCs for the three antibiotics by the DeLong test.

| Comparison | AUC□ | AUC□ | ΔAUC | Z statistic | P value |
| --- | --- | --- | --- | --- | --- |
| Amoxicillin vs levofloxacin hydrochloride | 0.949 | 0.941 | 0.008 | 0.121 | 0.904 |
| Amoxicillin vs clarithromycin | 0.949 | 0.774 | 0.175 | 1.685 | 0.092 |
| Levofloxacin hydrochloride vs clarithromycin | 0.941 | 0.774 | 0.167 | 1.603 | 0.109 |

Note: nonparametric DeLong method (DeLong ER et al., Biometrics, 1988); two-sided test, $\alpha = 0.05$ .

## 4. Discussion

### 4.1 Value and feasibility of F-ROSE for H. pylori susceptibility testing

Our data confirm that fluorescence detection combined with AI recognition shortens the time to a susceptibility report substantially, with results available within a few hours, so that testing at the time of endoscopy becomes a realistic prospect. Apart from fluorescence-based instruments, microfluidic colorimetric rapid AST systems can also complete testing within 4–6 h but are not adapted for gastric biopsy samples either^[12]^. While commercial fluorescence-based rapid AST platforms such as the Accelerate Pheno® System also deliver results in several hours, they are exclusively validated for blood culture pathogens and lack optimization for gastric H. pylori specimens and drug-induced coccoid morphotypes^[13]^. In the pooled analysis of both rounds, overall sensitivity across the three antibiotics was 96.0% and NPV 97.8%, which indicates that the method identifies susceptible isolates reliably. In clinical terms, when F-ROSE classifies a strain as susceptible to a particular agent, the physician can build an eradication regimen around that drug with reasonable confidence, and the risk of failure caused by inadvertent use of an antibiotic to which the strain is resistant falls considerably.

What clinical practice asks of a rapid assay is above all that resistant strains are not missed, and the combination of high sensitivity and high NPV observed here fits that requirement well.

### 4.2 Gains in diagnostic performance after algorithm optimization

The central finding of this study is that improving fluorescence image clarity and upgrading the AI recognition model for *H. pylori* together raised the diagnostic accuracy of the whole F-ROSE assay appreciably. Pooled accuracy across the three antibiotics rose from 77.8% to 81.0%; the gain was largest for clarithromycin, where agreement moved from fair to moderate, and agreement for amoxicillin improved as well.

That clarithromycin benefited most is plausibly related to the greater morphological heterogeneity of clarithromycin-resistant organisms. As systematically summarized by Krzyżek and Grande, clarithromycin stress strongly induces H. pylori to shift into morphologically diverse coccoid phenotypes with variable ultrastructural features, which creates substantial challenges for bacterial identification and susceptibility testing^[14]^. H. pylori converts more readily to coccoid variants under clarithromycin exposure, and the training set of the first-generation algorithm contained too few atypical forms for these to be recognized reliably; after optimization the expanded training set covered a range of variant morphologies and, combined with noise-reduction imaging, made missed identification of atypical organisms much less likely. Of note, the improved signal-to-noise ratio also made bacteria easier to distinguish from tissue debris and background fluorescence in fields of low bacterial density, which further reduced recognition bias.

### 4.3 Reasons for the low positive predictive value of F-ROSE and possible remedies

PPV was generally low across both rounds for all three antibiotics, with most estimates below 70%, indicating a certain proportion of false-positive readings. Taken together with the design and the statistical results of this study, three main contributors can be identified.

The first is the limited total sample and the small absolute number of resistant isolates. PPV depends heavily on the prevalence of resistance in the population studied, and among the 29 evaluable samples only 3 were amoxicillin-resistant, so that even a few false positives depress PPV markedly; with samples this small, the value obtained cannot be taken to reflect the true predictive performance in a clinical population. Consistent with this, the forest plot (Figure 3) shows very wide 95% CIs for the 14–15 samples available in each single round, so that statistical precision is poor and the uncertainty of the estimates correspondingly greater.

The second is that the uniform FRR cutoff of 50% has never been calibrated on a large sample. We adopted 50% as the boundary between susceptible and resistant on the basis of a small pilot experiment, without systematic threshold selection. ROC analysis of the full sample showed that the optimal cutoff for each of the three antibiotics lay above 50%: for levofloxacin hydrochloride, adjustment to the optimal FRR of 0.5812 raised specificity from 71.4% to 90.5% and PPV from 57.1% to 80.0%, with a substantial reduction in false positives, and for amoxicillin raising the cutoff to 0.5556 improved specificity modestly. Drug-specific FRR thresholds therefore appear capable of cutting the false-positive rate while preserving 100% sensitivity and, with it, the assurance that resistant strains are not missed.

The third is that non-specific background fluorescence cannot yet be eliminated entirely. Fragments of gastric mucosal tissue bind the fluorescent probes non-specifically and are easily read by the AI model as signal from live bacteria, which inflates the calculated FRR in antibiotic-treated wells and leads to susceptible strains being reported as resistant. A debris-recognition module added during algorithm optimization now separates *H. pylori* from mucosal fragments to a first approximation, but filtering of complex background fluorescence still leaves room for improvement. Refining the target specificity of the fluorescent probes and introducing image masking algorithms for debris are among the options for reducing interpretive bias from non-specific background signal further.

### 4.4 Limitations

This was a single-center, small-sample, prospective sequential diagnostic study, and both design and sample size impose several constraints on the generalizability of the findings and on clinical uptake.

First, the evaluable sample was small and statistical reliability correspondingly limited. Only 29 specimens grew successfully across the two rounds, with 14–15 evaluable samples per round; all single-round estimates in the forest plot carry wide 95% CIs and the point estimates fluctuate considerably, so that no firm conclusion about diagnostic performance can rest on a single round. Even after pooling, when the intervals narrowed, the total sample remains too small for precise estimation of the true value of each measure.

Second, culture failure introduces potential selection bias. Culture failed in 11 of the 40 enrolled patients because of bacterial death, insufficient bacterial load or contamination, and these cases were excluded outright. Since no reference-standard susceptibility result was available for them, they could not enter the analysis, and their absence may have inflated or deflated the apparent diagnostic performance of F-ROSE.

Third, statistical power was insufficient to settle the between-drug comparisons. All pairwise DeLong comparisons of the three AUCs gave P values above 0.05. This does not mean that discrimination is equivalent across the three antibiotics; rather, the sample was too small for the test to detect a real difference.

Fourth, the range of drugs tested was narrow. Only the three first-line agents for *H. pylori* eradication — amoxicillin, levofloxacin and clarithromycin — were included, and commonly used second-line drugs such as metronidazole and tetracycline were not, so the performance of the assay for these agents remains unknown.

Fifth, a single-center design limits the range of conditions sampled. All specimens came from one center, with uniform endoscopic technique, specimen transport and laboratory incubation conditions. Equipment and workflows differ between institutions, so the present conclusions cannot be transferred directly to a multicenter clinical setting and require validation in external cohorts.

## 5. Conclusion

Through two rounds of paired testing, this study assessed the diagnostic performance of F-ROSE in *H. pylori* susceptibility testing and the effect of algorithm optimization. Three points emerge. (1) F-ROSE offers high sensitivity and NPV for *H. pylori* susceptibility testing and is capable of rapid screening for resistance in clinical practice, which can help guide the choice of antibiotics. (2) After fluorescence image acquisition was made clearer and the AI recognition algorithm improved, overall diagnostic performance in round 2 was clearly better than in round 1: sensitivity and NPV both reached 100.0%, accuracy rose to 81.0%, false negatives fell to zero, and the gain was greatest for clarithromycin. (3) PPV remains low, indicating that there is still scope for reducing false positives.

Taken together, the optimized F-ROSE assay shows good accuracy and clear clinical potential for rapid *H. pylori* susceptibility testing, with a particularly strong performance in reducing missed detection. Future work should concentrate on enlarging the sample, refining the FRR decision threshold, extending the panel of drugs tested and undertaking multicenter validation, so as to move the technique toward routine clinical use.

